# Paternal obesity and epigenetic inheritance of breast cancer: The role of systemic effects and transmission to the second generation

**DOI:** 10.1101/2020.06.05.136234

**Authors:** Camile C. Fontelles, Raquel Santana da Cruz, Alexandra K. Gonsiewski, Ersilia Barin, Volkan Tekmen, Lu Jin, M. Idalia Cruz, Olivier Loudig, Anni Warri, Sonia de Assis

## Abstract

**Background:** While genetics explains some familial breast cancer cases, we showed that environmentally-induced epigenetic inheritance of breast cancer can also occur in rodent models. We previously reported that paternal consumption of a high-fat diet and ensuing obesity increased breast cancer susceptibility in the offspring (F1). Nevertheless, it is still unclear whether paternal-induced programming of breast cancer in daughters is associated with systemic alterations or mammary epithelium-specific factors. It also remains to be determined whether the ancestrally programmed breast cancer predisposition in F1 progeny can be transmitted to subsequent generations.

**Methods:** Male mice (F0) were fed either a control (CO) diet or an obesity-inducing diet (OID) for seven weeks and then mated with female mice (F0) reared on a CO diet. The resulting offspring (F1), also exclusively fed CO diet, were either used for mammary gland and tumor transplantation surgeries or to generate the F2 generation. To induce the mammary tumors, female mice were treated with 7,12 dimethylbenz[a]anthracene (DMBA). Total RNA extracted from F0 or F1 males sperm was used for small RNA-Seq analysis.

**Results:** Mammary glands from F1 CO female offspring exhibited enhanced development when transplanted into OID females [OID(CO-MG)], as shown by higher mammary gland area, epithelial branching and elongation, compared to CO females that received a CO mammary gland [CO(CO-MG)]. Similarly, mammary tumors from F1 CO female offspring transplanted into OID females [OID(CO.T)] displayed improved growth with a higher proliferation/apoptosis rate. We also found that granddaughters (F2) from the OID grand-paternal germline showed accelerated tumor growth compared to COxCO granddaughters (F2). Transmission of breast cancer predisposition to the F2 generation through OID male germline was associated with alterations in specific sperm tRNA fragments (tRF) in both F0 and F1 males.

**Conclusions:** Our findings indicate that systemic metabolic and mammary stromal alterations are the most significant contributors to paternal programming of mammary gland development and cancer predisposition in female offspring rather than mammary epithelium confined factors. Our data also show breast cancer predisposition in OID daughters can be transmitted to subsequent generations and could explain some familial cancers, if confirmed in humans.

## Introduction

Genetic predisposition explains most but not all familial diseases, including breast cancer[1]. It is increasingly evident that epigenetic inheritance of disease can also occur and may explain some inherited conditions. There is strong indication that, at conception, parents pass more than genetic material to their offspring. They also transmit a molecular memory of past environmental exposures [2, 3] which can result in offspring’s predisposition for certain chronic diseases [4].

Life-style and environmental insults have been shown to reprogram the sperm epigenome in humans and in animal models [5, 6]. Recently published studies demonstrated that the small RNA load in paternal sperm can convey phenotypes to the progeny [3, 7–9]. Some of those reports implicate t-RNA fragments (tRFs)— which are the most abundant small RNA sub-type in sperm—in the transmission environmentally-induced information from fathers to offspring and show that they can recapitulate disease phenotypes [7–10].

Because mammary gland development starts during fetal development, multiple studies report that maternal exposure during gestation can epigenetically reprogram the daughters’ mammary tissue and increase breast cancer development [11–14]. However, a role for paternal exposures in modulating breast cancer predisposition in offspring has emerged in recent years. We recently showed that paternal obesity, malnutrition and consumption of a high-fat diet all lead to increased breast cancer development in offspring [15–17], a phenotype associated with changes in normal mammary gland development. We also found that a recurrent phenotype accompanying offspring’s cancer predisposition is metabolic dysfunction [16–18], raising the possibility that paternally-induced cancer development could be a function of both systemic effects as well as tissue specific changes.

Paternal effects on the F1 generation include alterations in the germline epigenome [19], suggesting that disease traits in offspring could be passed on to future generations. Indeed, it has been reported that paternally-induced phenotypes observed in the F1 can be transmitted to the F2 generation [19, 20]. It is still not clear, however, whether paternally-induced breast cancer predisposition observed in the offspring can be transmitted through successive generations without continuous exposure to the initial insult.

Here, we used a mouse model of paternal obesity and aimed to address the role of systemic metabolic alterations and the local mammary epithelial and/or stromal changes on breast cancer development in the F1 generation. We also investigated whether the breast cancer predisposition observed in daughters of obese fathers could be transmitted to granddaughters.

## Material and Methods

### Dietary Exposures and Breeding

The C57BL/6 mouse strain was used in all experiments. Male mice were fed AIN93G-based diets containing either 17.2 % (Control, CO, Envigo-Teklad #TD160018) or 57.1% (Lard-based, Obesity-Inducing-Diet, OID, Envigo-Teklad #TD160019) energy from fat (Diet details in supplementary **Table S1**, see the section on supplementary data) starting after weaning (3 weeks of age). Males’ body weight was recorded weekly (**Fig. S1**). At 10 weeks of age, OID-fed and CO-fed F0 male mice were mated with female mice reared solely on the CO diet to generate the F1 generation. Males were kept in female cages for 3 days. Female mice were kept on the CO diet during the breeding period, for the extent of pregnancy (21 days) and after giving birth. The birth weight and number of pups per litter were determined 2 days after birth. To avoid litter-effect, pups were cross-fostered one day after dams gave birth. Pups from 2–3 dams were pooled and housed in a litter of 8—10 pups per nursing dam. All pups were weaned on postnatal day 21 and fed the CO diet throughout the experiment. Pups body weight was recorded weekly.

To obtain the F2 generation, F1 male offspring from OID fathers were mated with F1 females from either CO [OIDxCO] or OID [OIDxOID] groups. Similarly, F1 male offspring from CO fathers were mated with F1 females from either the CO [COxCO)] or OID [COxOID] groups. No sibling mating was carried out. F1 and F2 generation females from the CO or OID lineages were used to study body weight, metabolic function, mammary tumorigenesis and mammary transplantation, as described in the following sections. The experimental design is shown in **Fig. S2**.

<u>F1 and F2 litters’ gender distribution and number of offspring used in each experiment are shown in **Table S2** and **Table S3**, respectively</u>. All animal procedures were approved by the Georgetown University Animal Care and Use Committee, and the experiments were performed following the National Institutes of Health guidelines for the proper and humane use of animals in biomedical research.

### Metabolic Function

Insulin tolerance test (ITT) was performed after the mice fasted for 6 h, according to the method described by Takada et al [21]. The insulin load (75 mU/100 g body weight) was injected as a bolus, and the blood glucose levels were determined at 0, 3, 6, 9, 12, and 30 minutes after injection in female offspring. The area under the curve (AUC) was calculated according to the trapezoid rule. Differences in ITT were analyzed using two-way ANOVA (group, time), followed by post-hoc analyses.

### Mammary Transplantation

Three-week old F1 female offspring of CO and OID males underwent a mammary gland transplantation surgery as previously described [22, 23]. The experimental design is shown in Fig. S3. Females undergoing surgery were anesthetized using isoflurane flowing in oxygen and maintained with isoflurane flowing at 1-3%. Before transplantation, the 4^th^ inguinal mammary gland of **host** females was cleared from their endogenous epithelium by removing the fat pad of the 4^th^ gland up to its proximal lymph node. Special care was taken to cut off the connection between the 4^th^ and 5^th^ mammary glands to ensure complete clearing of the 4^th^ mammary fat pad and to avoid later epithelial contamination from the 5^th^ mammary gland. The excised fat pad containing the epithelial cells were stained with carmine aluminum solution to check cleared margins.

For transplantation, the **donor** fat pad containing the epithelial cells was excised and divided into small pieces (1mm^3^) and placed into a tissue-culture plate containing DMEM/F12 to keep it moist. Mammary tissue fragments of the donor mouse, either CO or OID F1 female offspring, were then implanted into a pocket made in the cleared fat pad of the host (CO or OID). The skin incision was closed with surgical wound clips. The transplantations were performed from CO female offspring donors to both CO [CO(CO-MG)] and OID [OID(CO-MG)] female offspring hosts, as well as from OID female offspring donors to CO [CO(OID-MG)] female offspring hosts. Mammary glands transplants were collected approximately 10 weeks post-surgery and used for analysis of epithelial branching density, epithelial elongation and number of Terminal End Buds (TEBs) as described in the next sections.

### Transplanted mammary gland growth and development

Transplanted mammary glands collected approximately 10 weeks post-surgery were stretched onto a slide, placed in a fixative solution and stained with a carmine aluminum solution (Sigma Chemical Co.) as previously described [24]. Whole mounts were examined under the microscope (AmScope) for ductal elongation and number of TEBs (undifferentiated structure considered to be the targets of malignant transformation), as previously described [24]. Whole-mount slides were also photographed (Olympus SZX12 250 Stereomicroscope), digitized and analyzed. Briefly, the portion surrounding the glandular epithelium was removed, color channels separated, and noise removed. The images were thresholded and skeletonized. Then, mammary epithelial area and branching (sum of intersections) were measured by Sholl analysis, a plugin ImageJ software (National Institute of Health, Bethesda, MD, USA) as previously described [25]. Once morphological analyses were completed, mammary whole mounts were removed from the slide, embedded in paraffin, sectioned (5 μm) [26] and prepared for either hematoxylin and eosin (H&E) or ki-67 staining as described below. Differences between groups were analyzed using one-way ANOVA, followed by post-hoc analyses.

### Mammary tumor induction

Mammary tumors were induced in F1 an F2 female offspring by administration of medroxyprogesterone acetate (MPA; 15 mg/100μl, subcutaneously) to 6 weeks of age female offspring, followed by three weekly doses of 1 mg 7,12-dimethylbenz[a] anthracene (DMBA; Sigma, St. Louis, MO) dissolved in peanut oil by oral gavage[27]. Tumors were detected by palpation once per week, starting at week 2 after the last dose of DMBA. Tumor growth was measured using a caliper, and the width and height of each tumor were recorded.

In the F1 generation, mammary tumors were harvested when reaching approximately 40 mm^2^ in volume and used for mammary tumor transplantation surgery, as described in the next section. In the F2 generation, tumor development was monitored for a total of 20 weeks post-DMBA administrations. Animals in which tumor burden reached approximated 10% of total body weight were euthanized before the end of the monitoring period, as required by our institution. Tumor growth was analyzed using two-way ANOVA (group and time), followed by post-hoc analyses. Kaplan-Meier survival curves were used to compare differences in tumor incidence, followed by the log-rank test. Differences in tumor latency and mortality were analyzed using two-way ANOVA.

### Mammary tumor transplantation

CO and OID F1 female offspring underwent a mammary tumor transplantation surgery at approximately 11 weeks of age. Females undergoing surgery were anesthetized using isoflurane flowing in oxygen and maintained with isoflurane flowing at 1-3%. Briefly, carcinogen-induced mammary tumor fragments (1 mm^3^) of a donor mouse, either CO or OID offspring, were implanted into a pocket made in the mammary fat pad of the host (CO or OID). The experimental design is shown in Figure S3. Mammary tumors grown from the transplants were collected approximately 6-8 weeks post-surgery. Differences between groups were analyzed using one-way ANOVA, followed by post-hoc analyses.

### Analysis of cell proliferation

Cell proliferation (Ki-67) was evaluated by immunohistochemistry in mammary gland and mammary tumors transplants. Briefly, tissues were fixed in 10% buffered formalin, embedded in paraffin, and sectioned (5 μm). Sections were deparaffinized with xylene and rehydrated through a graded alcohol series. Antigen retrieval was performed by immersing the tissue sections at 98°C for 40 minutes in 1X Diva Decloaker (Biocare). Tissue sections were treated with 3% hydrogen peroxide and 10% normal goat serum for 10 minutes and were incubated with the primary antibody, overnight at 4°C. After several washes, sections were treated to the appropriate HRP labeled polymer for 30 min and DAB chromagen (Dako) for 5 minutes. Slides were counterstained with Hematoxylin (Fisher, Harris Modified Hematoxylin), blued in 1% ammonium hydroxide, dehydrated, and mounted with Acrymount. The sections were photographed using an Olympus IX-71 Inverted Epifluorescence microscope at 40x magnification. Proliferation index (Ki-67 staining) was determined by immunoRatio, a plugin Image J software (National Institute of Health, Bethesda, MD, USA), to quantify hematoxylin and DAB-stained cells. Differences between groups were analyzed using one-way ANOVA, followed by post-hoc analyses.

### Analysis of cell apoptosis

Cell apoptosis analysis was performed in transplanted mammary glands and tumors by morphological detection. Tissues were fixed in neutral buffered 10% formalin, embedded in paraffin, sectioned (5 μm) and stained with hematoxylin and eosin (H&E). Cells presenting loss of adhesion between adjacent cells, cytoplasmic condensation and formation of apoptotic bodies were considered apoptotic as described before[28]. Sections were photographed using an Olympus IX-71 Epifluorescence microscope at 40x magnification. Twenty areas were photographed randomly, and the number of apoptotic bodies counted. Images were evaluated with ImageJ software (NIH, USA). Differences between groups were analyzed using one-way ANOVA, followed by post-hoc analyses.

### Mature spermatozoa collection and purification

CO and OID-fed males (F0) and their male offspring (F1) were euthanized and their caudal epididymis dissected for sperm collection. The epididymis was collected, punctured, and transferred to tissue culture dish containing M2 media (M2 Medium-with HEPES, without penicillin and streptomycin, liquid, sterile-filtered, suitable for mouse embryo, SIGMA, product #M7167) where it was incubated for 1 hour at 37°C. Sperm samples were isolated and purified from somatic cells. Briefly, the samples were washed with PBS, and then incubated with SCLB (somatic cell lysis buffer, 0.1% SDS, 0.5% TX-100 in Diethylpyrocarbonate water) for 1 hour. SCLB was rinsed off with 2 washes of PBS and the somatic cell-free purified spermatozoa sample pelleted and used for RNA extraction.

### Small RNA-Seq and Gene Ontology (GO) analyses

Total RNA was isolated from sperm using Qiagen’s miRNeasy extraction kit, according to the manufacturer’s instructions. One hundred ng of column-purified sperm RNA was used to prepare individually barcoded small-RNA libraries. Samples were barcoded, pooled, precipitated and separated on a 15% polyacrylamide gel (PAGE). The gel was stained with SYBR^®^ gold dye and the small non-coding RNA segment corresponding to transfer RNA fragments or tRFs (30-45 nucleotides) excised and purified using a cDNA library preparation method described previously [29]. This library preparation method was demonstrated to be highly reproducible using total RNA with RNA Integrity Numbers as low as 2.0[29]. Indexed, single-ended small-RNA sequencing libraries were prepared. For each individual barcoded library, at least 10 million reads (raw data) were generated using an Illumina Hi-Seq 2500. The raw reads were subjected to 3’ adapter trimming and low quality filtering using Trimmomatic program [30]. The high quality clean reads were aligned to the mouse genome. tRFs tags were mapped to the mouse genome (GRCm38/mm10 reference genome) in order to analyze their genomic distribution and expression in the different sperm RNA samples. Small RNA tags were annotated and aligned to known t-RNA sequences using Ref-seq, GenBank and Rfam database using blastn with standard parameters. To analyze the differential expression of tRFs between CO and OID groups, tRFs were normalized to TPM (Transcripts Per Kilobase Million). tRFs with a P value less than 0.05 were considered significant, with an appropriate correction for multiple testing [31]. Target genes for the 5 overlapping tRFs in OID F0 and F1 males were predicted using TargetScan Mouse custom seedmatch and modified miRanda algorithm (energy <= −20 and score >= 150). The common predicted genes were then uploaded to PANTHER 15.0 for GO term and pathway analysis, final lists were filtered by FDR < 0.25.

## Results

### Offspring of OID fathers have impaired metabolic function and altered mammary gland development

We previously reported that paternal consumption of obesity-inducing diets (OID) at the preconception window increased female offspring’s susceptibility to breast cancer [15, 16]. In those studies, we also described mammary gland morphological changes as well as metabolic dysfunction—a phenotype also reported by others—in offspring of obese fathers [16, 18, 19, 32]. Our present results corroborate our previous findings as OID offspring (F1) displayed impaired metabolic function with both F1 males and females showing significantly reduced insulin sensitivity compared to CO offspring (P=0.002, P=0.011, **Fig. 1a-f**). In addition, mammary glands of OID daughters also showed increased number of terminal end buds (TEB), higher epithelial branching and elongation, although only the last parameter reached statistical significance compared to CO (**Table S4**). Those phenotypes were not associated with body weight gain (**Fig. S4**) as OID offspring weights either did not differ from or were lower than CO.

**Figure 1:**
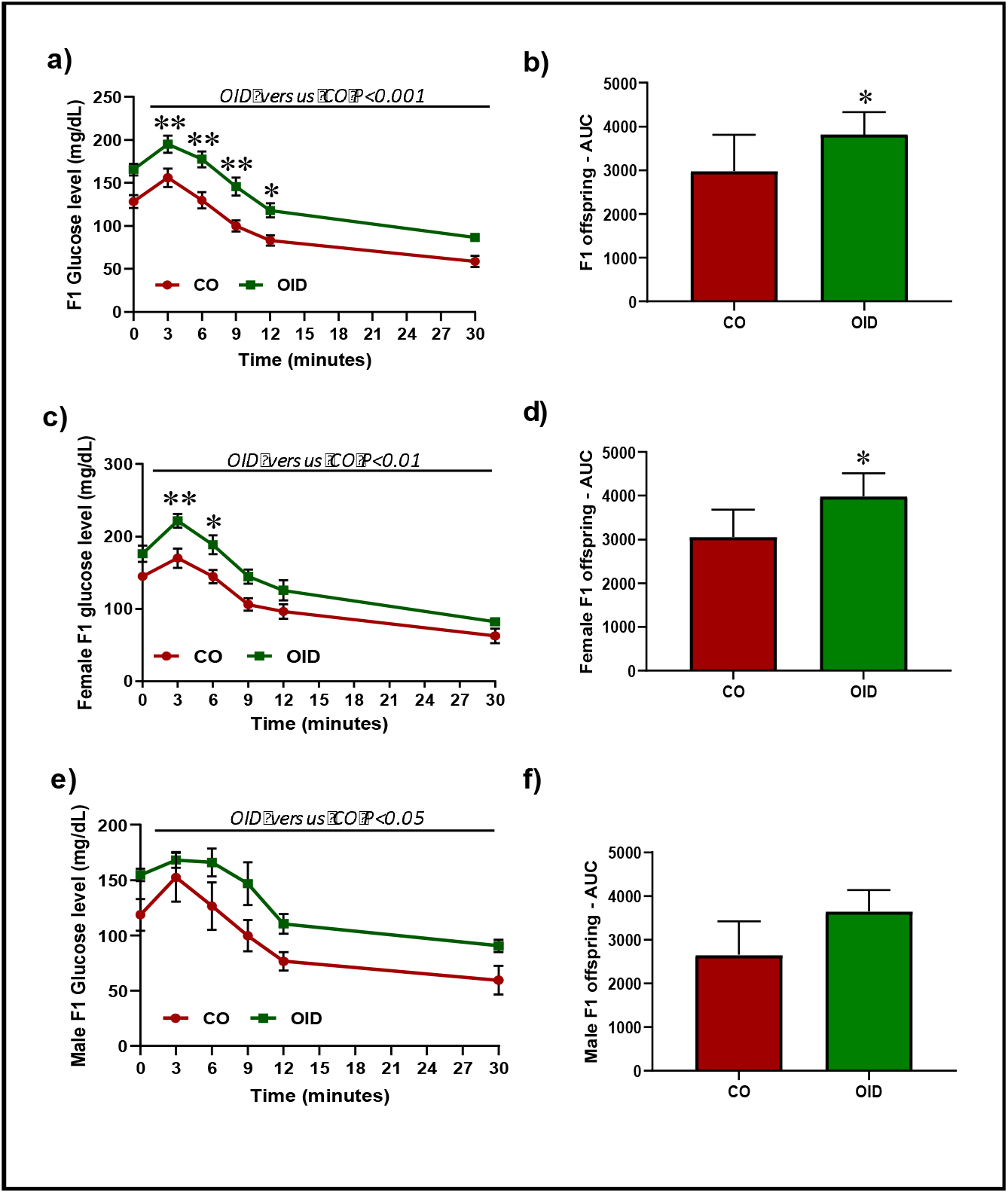
Paternal OID causes metabolic disturbance in offspring. Insulin tolerance test (ITT) and area under curve (AUC) in all gender (**a-b**), female (**c-d**) and male (**e-f**) F1 offspring (n=7-8/gender/group) from CO and OID-fed fathers. The data are expressed as mean ± SEM. Significant differences versus the control group were determined by two-way ANOVA followed by post-hoc analysis. *P≤0.05; **P≤0.01.

### Systemic effects play a larger role in normal mammary tissue and mammary tumor growth in offspring of OID fathers

Next, we examined the contributions of systemic alterations and mammary tissue specific factors (stroma *vs.* epithelium) to the increased breast cancer development in offspring of obese fathers. In the first experiment, female offspring of either CO or OID-fed males underwent a mammary gland transplantation surgery. CO mammary glands transplanted into OID females [OID(CO.MG)] exhibited accelerated development (**Fig. 2a-e**) as shown by higher mammary gland area (p=0.032, **Fig 2b**), higher mammary branching and higher epithelial elongation (p=0.014; p=0.008, respectively, **Fig. 2c-d**), but not higher number of TEBs (**Fig. 2e**), compared to CO females that received a CO mammary gland [CO(CO.MG)]. This phenotype was associated with a higher proliferation index and lower apoptotic rates compared to compared to [CO(CO.MG)]) and [CO(OID.MG)] (P=0.021 and P=0.026, respectively; **Fig. 2f-j**). While OID mammary glands transplanted into CO females [CO(OID.MG)] showed slightly higher mammary gland area, mammary branching and epithelial elongation and number of TEBS (**Fig.2b-e**) compared to [CO(CO.MG)], results did not reach statistical significance.

**Figure 2:**
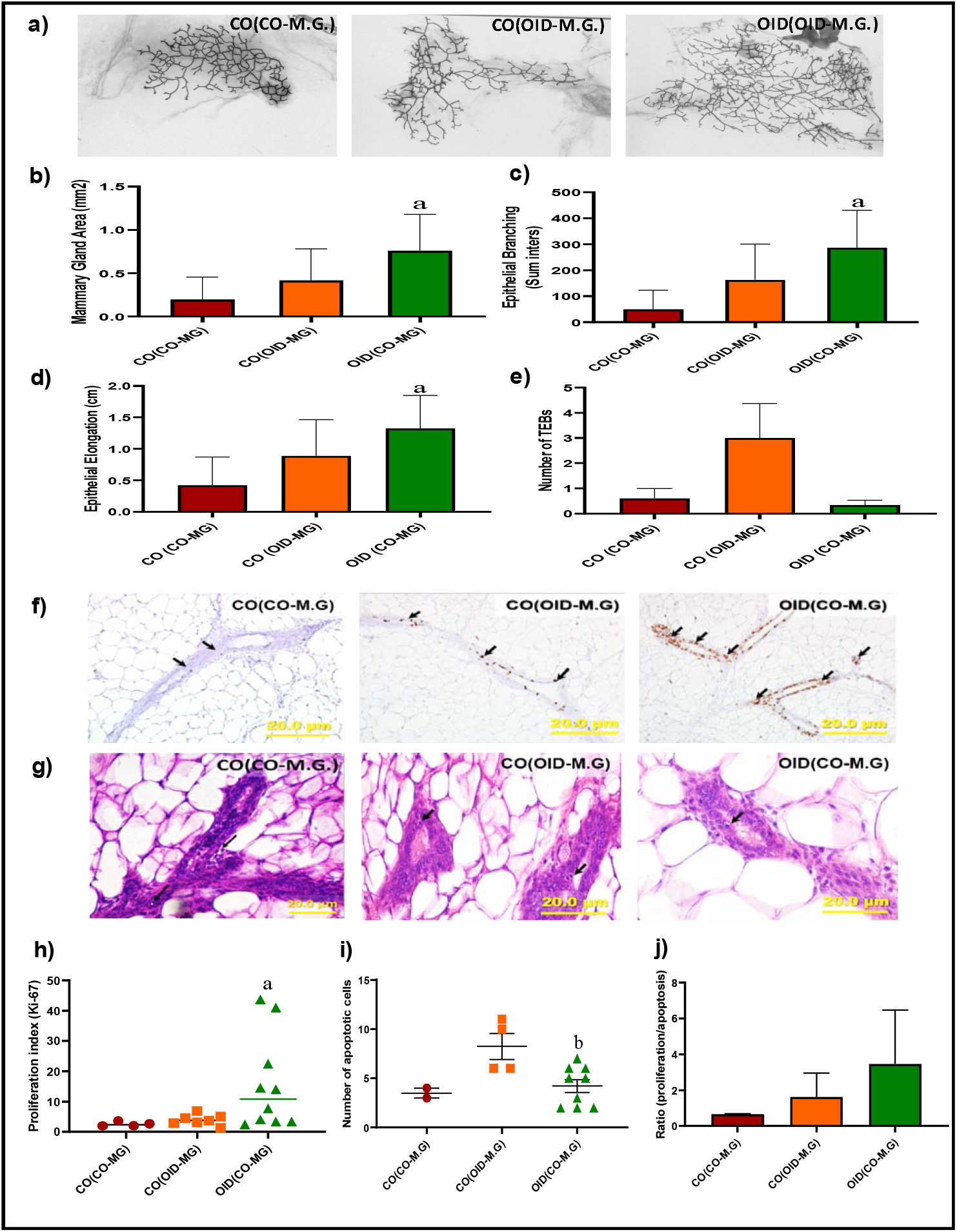
Development of transplanted mammary glands in CO or OID daughters (F1). Histological depiction of transplanted mammary gland in (**a**) [CO(CO-M.G)], [CO(OID-M.G)], and [OID(CO-M.G)] groups. Graphs below show values for mammary gland area (**b**), epithelial branching (**c)**, epithelial elongation (**d**) and number of terminal end buds (TEB) (**e**), (**b-e**, n=6-13); Photomicrograph of Ki-67 immunostaining (**f**) (20x, staining indicated by arrows) and apoptotic cells (**g**) (H&E morphological assessment, 40x, cells indicated by arrows). Graphs below show proliferation index (**h**), number of apoptotic cells (**i**) and proliferation/apoptosis ratio (**j**), (**f-i**, n=4-12). The data are expressed as mean ± SEM. Significance differences between groups were determined by one-way ANOVA followed by post-hoc analysis (mammary gland area, branching density, epithelial elongation, number of TEBs, cell proliferation and apoptosis numbers). “a” indicates statistically significant difference (P≤ 0.05) between OID(CO-M.G) and CO(CO-M.G); “b” indicates statistically significant difference (P≤0.05) between OID(CO-M.G) and CO(OID-M.G).

Given that both the mammary microenvironment and systemic response could play a role in tumor progression, we also asked whether the metabolic-induced mammary stroma milieu could affect the growth potential of tumors. Thus, in our second experiment, a DMBA-induced mammary tumor of F1 female offspring from CO (donor) was transplanted into the fat pad of a CO or OID female offspring (host) and *vice versa.* Tumor growth was followed for 6-8 weeks post-surgery. Consistent with what we observed for mammary gland transplants, we found that CO tumors transplanted into OID females [OID(CO.T)] displayed improved growth (**Fig. 3a**) and shorter latency (**Fig. 3b**) compared to CO or OID tumors transplanted in CO females [CO(CO.T) and CO (OID.T)], although differences among the groups did not reach statistical significance. [OID(CO.T)] tumor also showed significantly increased cell proliferation to apoptosis ratio, compared to both [CO(CO.T)] and [CO (OID.T)] (p=0.043, P=0.032, respectively; **Fig. 3c-g**).

**Figure 3:**
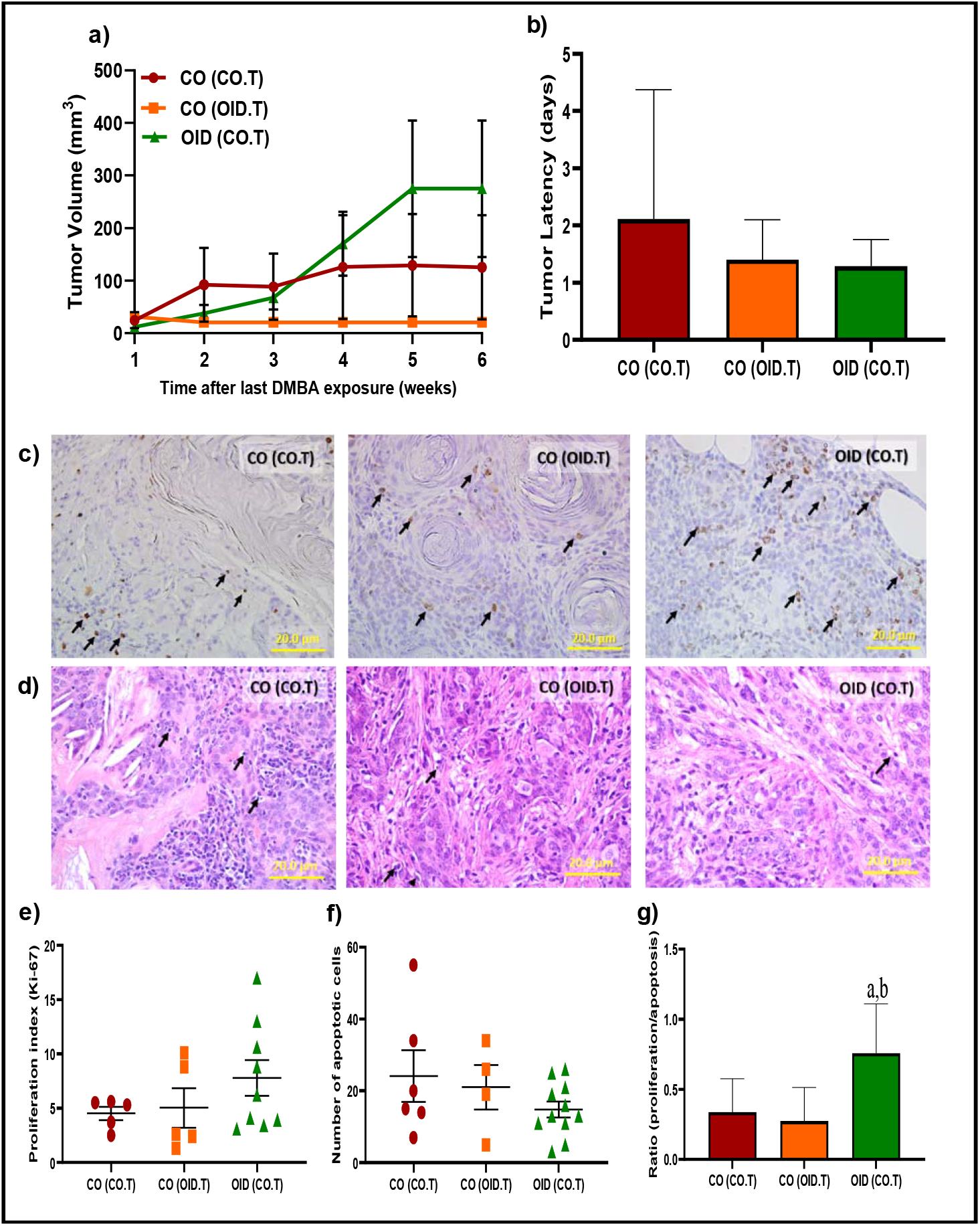
Development of transplanted mammary tumors in CO or OID daughters (F1). Tumor volume (**a**) and latency (**b**) (**a-b**, n=10-18/group) in [CO(CO-M.G)], [CO(OID-M.G)], and [OID(CO-M.G)] groups after a six-week monitoring period. Photomicrograph of Ki-67 immunostaining (**c**) (20x, staining indicated by arrows) and apoptotic cells (**d**) (H&E morphological assessment, 20x, cells indicated by arrows). Graphs below show proliferation index (**e**), number of apoptotic cells (**f**), and proliferation/apoptosis ratio (**g**), (**c-g**-n=3-11/group). The data are expressed as mean ± SEM. Significance differences between groups were analyzed by repeated measures ANOVA (mammary tumor volume) and one-way ANOVA (tumor latency, proliferation index and number of apoptotic cells) followed by post-hoc analysis. “a” indicates statistically significant difference (P≤0.05) between OID(CO.T) and CO(CO.T); “b” indicates statistically significant difference (P≤0.05) between OID(CO.T) and CO(OID.T).

### Consumption of OID alters the tRF content in sperm of fathers (F0) and their sons (F1)

Recent studies have suggested that sperm non-coding RNAs play a role in transmitting environmentally-induced information from fathers to offspring. Transfer RNA fragments or tRFs make up the majority of small RNAs in mature sperm and can recapitulate the effects of paternal obesity in offspring [3]. As reported before, GlyGCC and GlutCTC were the most abundant tRFs in sperm of both fathers (F0) and their male offspring (F1), representing about 70% of all tRFs (**Fig. 4a-b**) [8, 19]. We also found that consumption of OID altered specific tRFs in both father (**Fig. 4c**) and sons (**Fig. 4d**), with five tRFs overlapping between the two generations (**Fig. 4e**): Levels of ValTAC and SerCGA were increased while those of ArgCCG, ArgTCG and SeCTCA were decreased in sperm of OID F0 and F1 males compared to CO. Putative targets of these five tRFs were significantly enriched for molecular functions related to DNA binding, transcription factor activity, transcriptional regulation, and transmembrane transporters among others (**Fig. 4f**).

**Figure 4:**
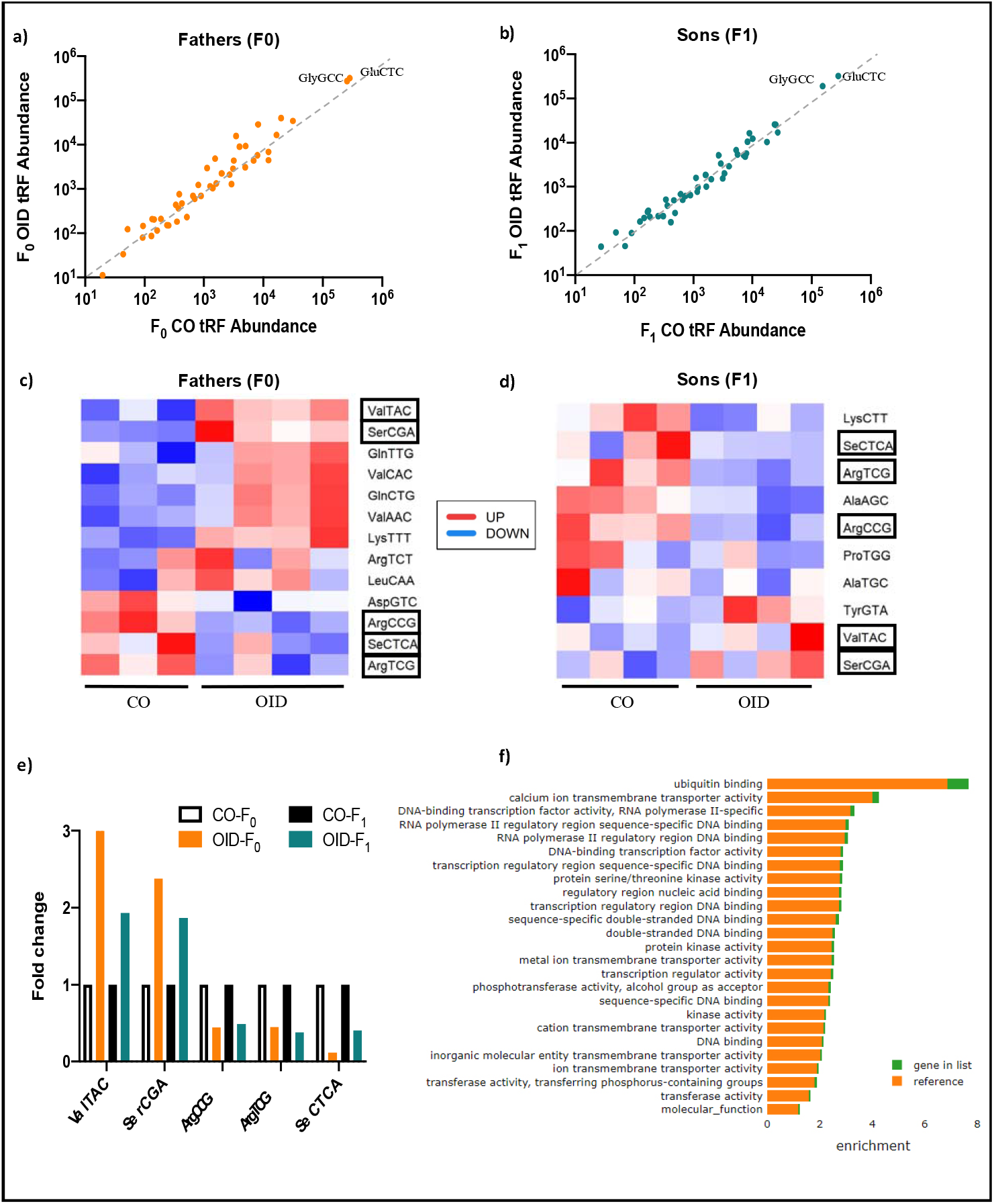
Paternal OID reprograms the sperm small non-coding RNA load in fathers (F0) and sons (F1). (**a-b**) Scatterplot of sperm tRNA fragments (tRF) from OID (y-axis) fathers (F0, **a**) and OID sons (F1, **b**) versus their respective controls (x-axis) (n=3-4/group) assessed by RNA-seq. GluCTC and GlyGCC are the dominant tRFs. (**c-d**) Heat-map showing differentially expressed tRNA fragments (tRFs) in sperm from OID fathers (**c**) and sons **(d**) compared to CO, highlighting overlapping tRFs in F0 and F1 (boxes). (**e**) Levels (fold change) of the 5 tRFs with overlapping differential expression in both OID fathers(F0) and sons (F1) compared to CO. (**f**) Gene ontology molecular functions significantly enriched in the targets of ValTAC, SerCGA, ArgCCG, ArgTCG and SeCTCA.

### Breast cancer predisposition in OID daughters is transmitted to a second generation

Given the tRF alterations observed in the F1 OID offspring germline, we then asked whether breast cancer predisposition in OID daughters could be inherited by a second generation of females. To this question, we produced the F2 generation by mating F1 male offspring from OID fathers with F1 females from either CO [OIDxCO] or OID [OIDxOID] groups. Similarly, F1 male offspring from CO fathers were mated with F1 females from either the CO [COxCO)] or OID [COxOID] groups (**Fig. S2**). Indeed, we found that the female F2 generation derived from either the F1 OID male and female lineage (OIDxCO and COxOID, respectively) or both (OIDxOID) developed carcinogen-induced mammary tumors that grew significantly faster, compared to COxCO group (p<0.001, **Fig.5a**). The incidence of mammary tumors at the end of the monitoring period was also significantly higher in F2 OIDxOID females compared to the COxCO group (p=0.037; **Fig. 5b**), suggesting a synergistic effect of both the male and female OID germlines. Tumor latency and tumor mortality rates in the OIDxCO group were slightly shorter than in all other groups, however results did not reach statistical significance (**Fig. 5c-d**). While all F2 females derived from the OID grand-paternal lineage (COxOID, OIDxCO, OIDxOID) showed higher mammary tumor growth with significantly larger tumors (**Fig. 5a**) when compared to COxCO, only OIDxOID females developed insulin insensitivity as shown by higher ITT and AUC values (p=0.007, P=0.017, **Fig. 5e-f**). However, OIDxCO females were significantly heavier overtime compared to all other groups (COxCO, COxOID, OIDxOID, p=0.0004, **Fig. S5**).

**Figure 5:**
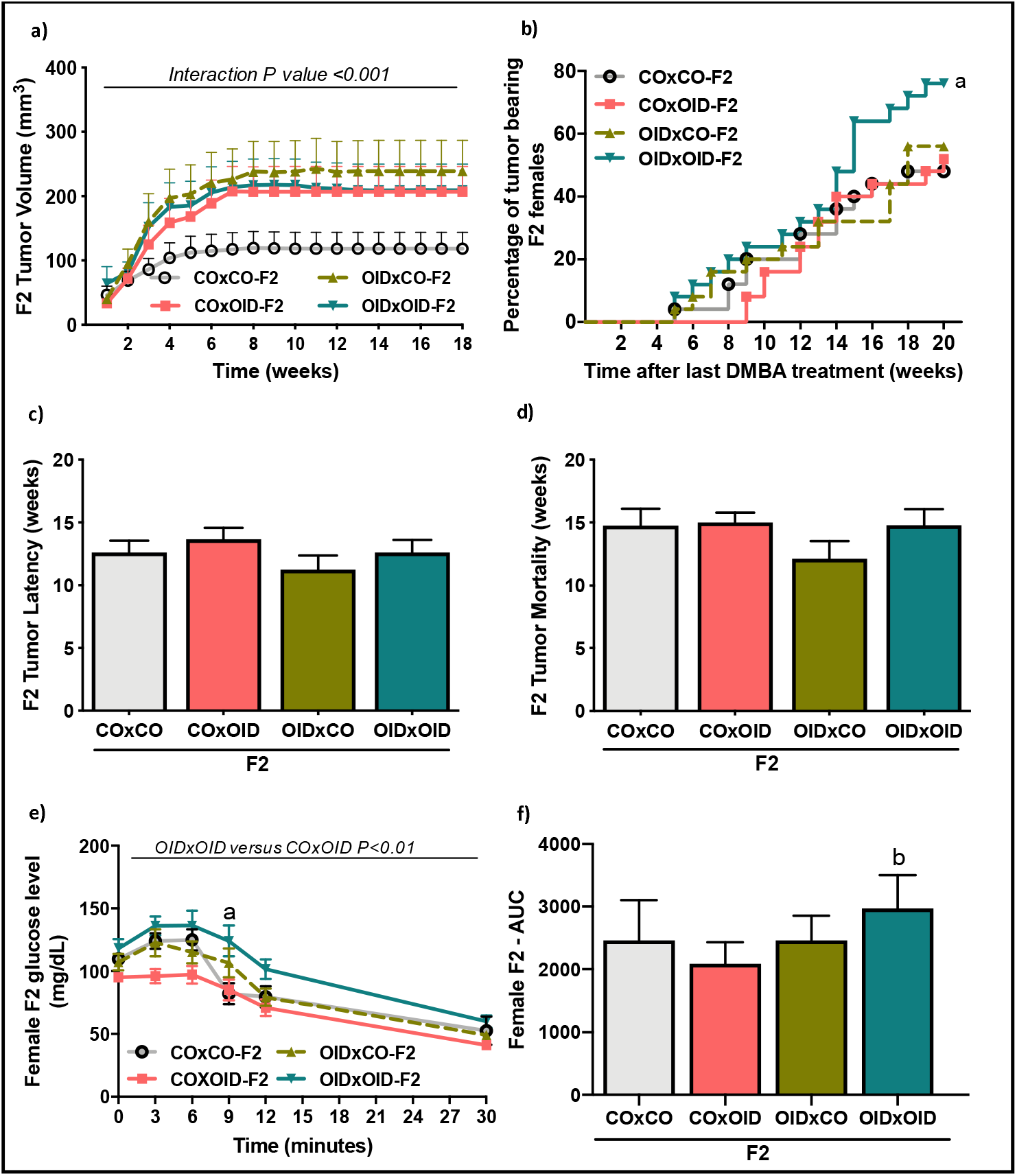
Paternal OID programs breast cancer development and metabolic dysfunction in granddaughters (F2). (**a-d**) Carcinogen-induced mammary tumorigenesis in CO and OID female **F2** offspring. Mammary tumor volume (**a**), tumor incidence (**b**), tumor latency (**c**) and tumor mortality (**d**) (n=25/group). Insulin tolerance test (ITT) (**e**) and (**f**) area under curve (AUC) in CO and OID female **F2** offspring (n=8/group). Tumor incidence is shown as percentage of animals with tumors. All other data are mean ± SEM. Significant difference were determined by Kaplan-Meier analysis followed by log-rank test (tumor incidence), repeated measures ANOVA (mammary tumor volume), one-way ANOVA (tumor latency, mortality and area under curve), or two-way ANOVA (ITT) followed by post-hoc analysis. “a” indicates statistically significant difference (P≤0.05) between OIDxOID and COxCO; “b” indicates statistically significant difference (P≤0.01) between OIDxOID and COxOID.

## Discussion

We previously reported that paternal obesity increases tumorigenesis in offspring, including breast cancer [15, 16, 18]. In this follow-up study, we showed that metabolic disturbances in the F1 generation play a key role in the increased breast cancer development observed in offspring of obese fathers in a mouse model. We also report that the paternal obesity leads to higher cancer development in two successive generations. Transmission of the increased breast cancer phenotype into the F2 generation was associated with epigenetic changes in the germline, namely alterations in the abundance of tRFs present in OID F1 male sperm.

The first aim of our study was to dissect the distinct contributions of systemic effects and mammary tissue-confined factors to increased breast cancer development in daughters of obese fathers, as we had observed both metabolic dysfunction and mammary gland abnormalities in previous studies[15, 16]. Our results showed that systemic metabolic effects, likely acting though the mammary stroma, in OID daughters play a larger role compared to the mammary epithelium. Further, tumors from CO offspring transplanted into OID daughters acquired a growth advantage compared to those transplanted in controls, suggesting that the stroma in OID females allows for better implantation and tumor growth. It is still possible that mammary epithelium confined factors play a role in the increased tumor development in OID offspring, however, they play a reduced role compared to systemic and mammary stromal effects according to our data. While it has been traditionally thought that the epithelium is the compartment with the dominant contribution regarding breast cancer initiation and growth and mammary tissue regeneration, some studies have highlighted the importance the stroma microenvironment, particularly adipocytes, on normal mammary development and malignant transformation of the mammary epithelium[33–36]. Our analyses are in agreement with those findings and suggest that the stroma plays an important enabling role for tumor growth.

It is also well established in epidemiologic studies that metabolic conditions such as obesity, metabolic syndrome and diabetes are important risk factors for breast cancer and other malignancies [37–40] and data from animal models offer support to those findings [41, 42]. In line with that, we demonstrated that a milieu of metabolic dysfunction and altered stromal microenvironment creates conditions for increased proliferation and survival of both normal and tumorigenic mammary cells as demonstrated by our transplantation studies.

While we have not directly investigated the molecular mechanisms behind the findings reported here, it is known that metabolic dysfunction contributes to cancer development via extrinsic and tumor-intrinsic factors [43]. Metabolic-induced alterations in growth factors signaling, inflammation and the associated microenvironment, as well as changes in tumor metabolism itself are all major contributors to cell proliferation and cancer development [43]. Not surprisingly, our previously reported results show that paternal obesity or malnutrition alters the molecular make-up of tumors which show increased growth factor and energy sensing signaling and altered amino-acid metabolism[15–18].

We also examined whether the offspring’s breast cancer predisposition programmed by paternal obesity could be inherited by a second unexposed generation. We found that the risk of breast cancer is passed down to the OID grandchildren equally via the F1 male and female germlines. Our data also suggest that there is a synergistic effect when both F1 parents had an obese father, with their descendants showing not only accelerated tumor growth but also higher tumor incidence. As with the F1 generation, F2 females from the OID lineage showed signs of metabolic dysfunction which depended whether they originated from the male or female lineage or both.

Our study offers some insights into the potential mechanism of transmission of breast cancer risk from one generation to another. Given the increased mammary tumorigenesis in the granddaughters of OID males in the absence of any further exposure, transmission of this phenotype conceivably occurs via F1 germ cells, which give rise to the F2 generation. In support of that, we found that F1 male germline showed alteration in tRFs, a class of small non-coding RNAs abundant in sperm, recently shown to transmit environmentally-induced information from one generation to another [7, 8]. While details on the functional role of tRFs in embryonic development are still under investigation, these small RNAs have been implicated in the regulation of translation, stress granule formation, viral replication and retrotransposons[44, 45]. Unfortunately, the inherent technical challenge of collecting enough eggs for molecular analysis precluded us from evaluating the F1 female germline. However, given that both the F1 male and female OID germline were able to transmit the increased predisposition to breast cancer phenotype to a second generation it is likely that we would have observed changes in the female germline as well. Nevertheless, we cannot rule out that some of the effects observed in F2 generation are due to maternal metabolic dysfunction in pregnancies of F1 OID females.

Interestingly, we found overlap in tRFs altered in sperm of F1 and F0 males. This suggests either that the F1 male germline is programmed by paternal obesity or that sperm non-coding RNAs are re-set in the F1 generation. Although, no changes in body weight were detected in F1 OID males, they did show metabolic dysfunction (impaired insulin sensitivity) later in life. However, others have shown that changes in the germline of male offspring of obese fathers occur in the absence of overt metabolic dysfunction[19], suggesting that F2 generation phenotypes represent true epigenetic inheritance.

The mechanisms for how germline epigenetic programming lead to phenotypes in offspring are still being investigated. However, given the short half-life of sperm small non-coding RNAs such as tRFs, is likely that they act early in embryonic development, setting a cascade of molecular events which biases cellular programming during subsequent divisions and culminate in disease phenotypes [3, 6]. Our gene ontology analysis of targets of the five overlapping tRFs in OID F0 and F1 OID males’ sperm showed an enrichment for functions related to DNA binding, transcription factor activity, transcriptional regulation, and transmembrane transporters. It is possible that an imbalance in the amount of those specific tRFs in sperm can disrupt embryonic development post-fertilization, programming the organism to be more to be more amenable and tolerant to cellular growth which would translate in increased cancer development. The exact mechanisms, however, need to be further investigated in a follow-up study.

In conclusion, the findings described here builds on our previous works and show that paternally-induced cancer development is largely due to systemic alterations in offspring and that the offspring’s breast cancer predisposition, as evaluated in this study, can be transmitted to a subsequent generation. While our study was conducted in an animal model, it could have important implications for human health. It is well known that family history is a strong predictor of cancer risk [46], yet not all familial cancers can be explained by genetic mutations[1, 47]. Though it is estimated the up to 30% of breast cancers cluster in families, only about one third of those are due to mutations in high penetrance genes such as *BRCA1* and *BRCA2,* leaving a sizable portion of familial breast cancers without a biological explanation [48]. Our study suggests that ancestral history of obesity from the paternal lineage could account for some familial cancers and that some organisms may be predisposed to the tolerance of cancer cells or may provide adequate conditions for their growth and development. This notion is supported by our prior findings showing that maternal exposure to an endocrine disruptor or dietary fat can also lead to multigenerational risk of breast cancer through both the male and female germlines in rats[12]. Given that the current study was performed in mice, our findings have now been confirmed in two different animal species.

It is also important to note that conditions such as obesity and malnutrition often occur in minorities and disadvantaged populations [49]. Our findings would suggest that social determinants of cancer predisposition and outcomes may be imprinted even before birth and are epigenetically mediated. However, it remains to be determined whether the biological insights uncovered by our study can account for some of the familial breast cancer predisposition or cancer disparities in humans.

## Acknowledgments

We thank the following Lombardi Cancer Center Shared Resources (SR) for their assistance: Animal Model SR, Histopathology & Tissue SR, Microscopy and Imaging SR. This study was supported by The American Cancer Society (RSG-16-203-01-NEC, Research Scholar Grant to S. de Assis), and the National Institutes of Health (1P30-CA51008; Lombardi Comprehensive Cancer Center Support Grant to Louis Weiner and CCSG pilot fund to S. de Assis).

The small RNA-seq data has been deposited in GEO (Gene Expression Omnibus) database with accession code pending.

## Author contributions

C.C.F, A.W. and S.D.A. conceived the study. S.DA. oversaw the research and wrote the manuscript with the help of C.C.F, R.S.C., A.W. and O.L.; C.C.F performed most of the experiments with the help of A.W., R.S.C., A.K., E.B. M.I.C. and V.T.; C.C.F. and R.S.C. analyzed the data; O.L. performed the sperm small RNA-seq profiling and L.J. performed the RNA-seq data and GO analysis.

## Supplementary Material

### Supplementary Figures Legends

**Figure S1:**
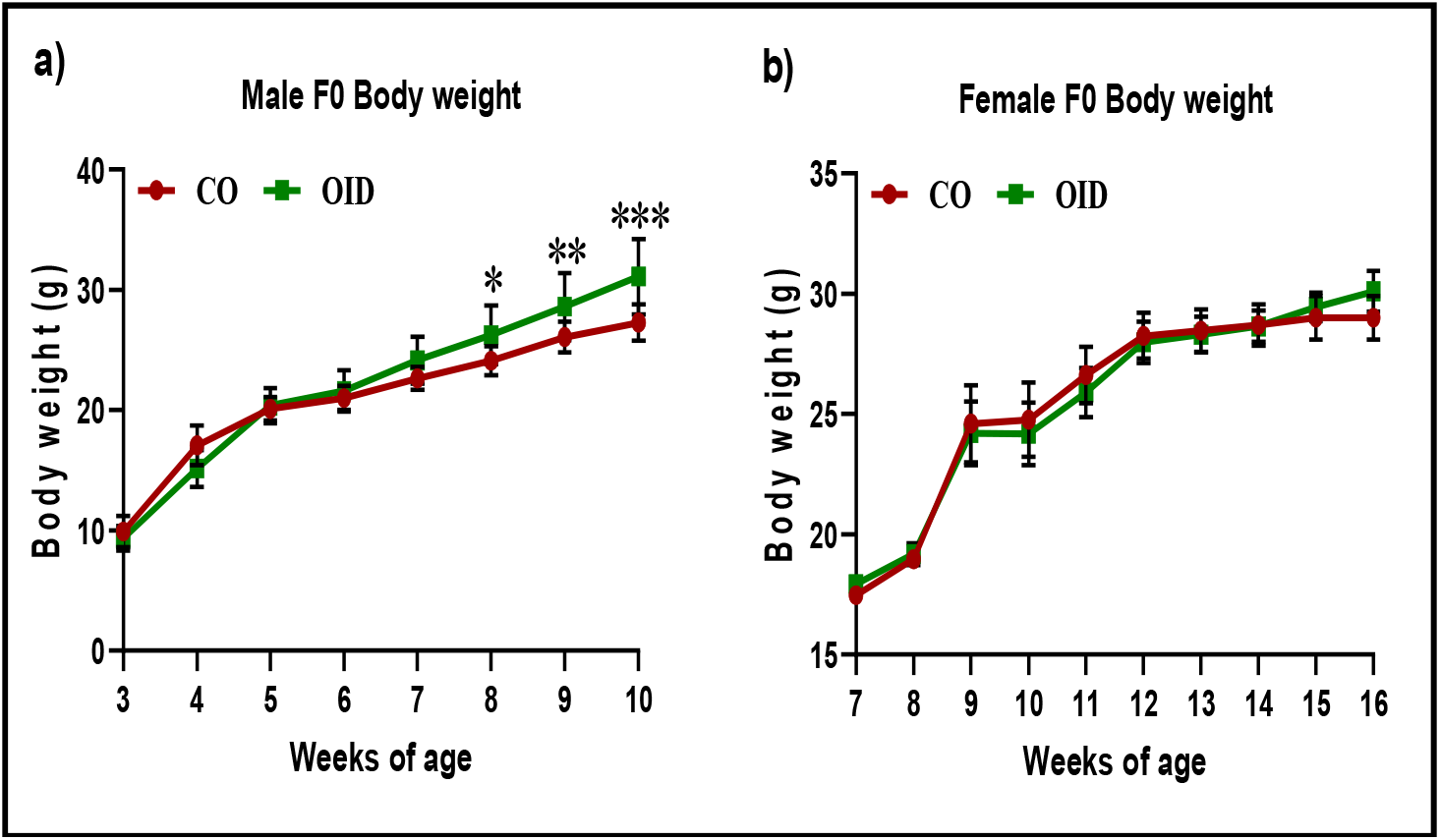
Parental body weight gain: **a**) Longitudinal body weight in control (CO, n=12) and obesity-inducing diet (OID, n=11) fed male mice sires. **b**) Longitudinal body weight of pregnant dams mated with CO (n=16) or OID (n=19) males. The data are expressed as mean ± SEM. Significant differences versus the control group were determined by two-way ANOVA followed by post-hoc analysis. *P≤0.05; **P≤0.01; ***P≤0.001

**Figure S2:**
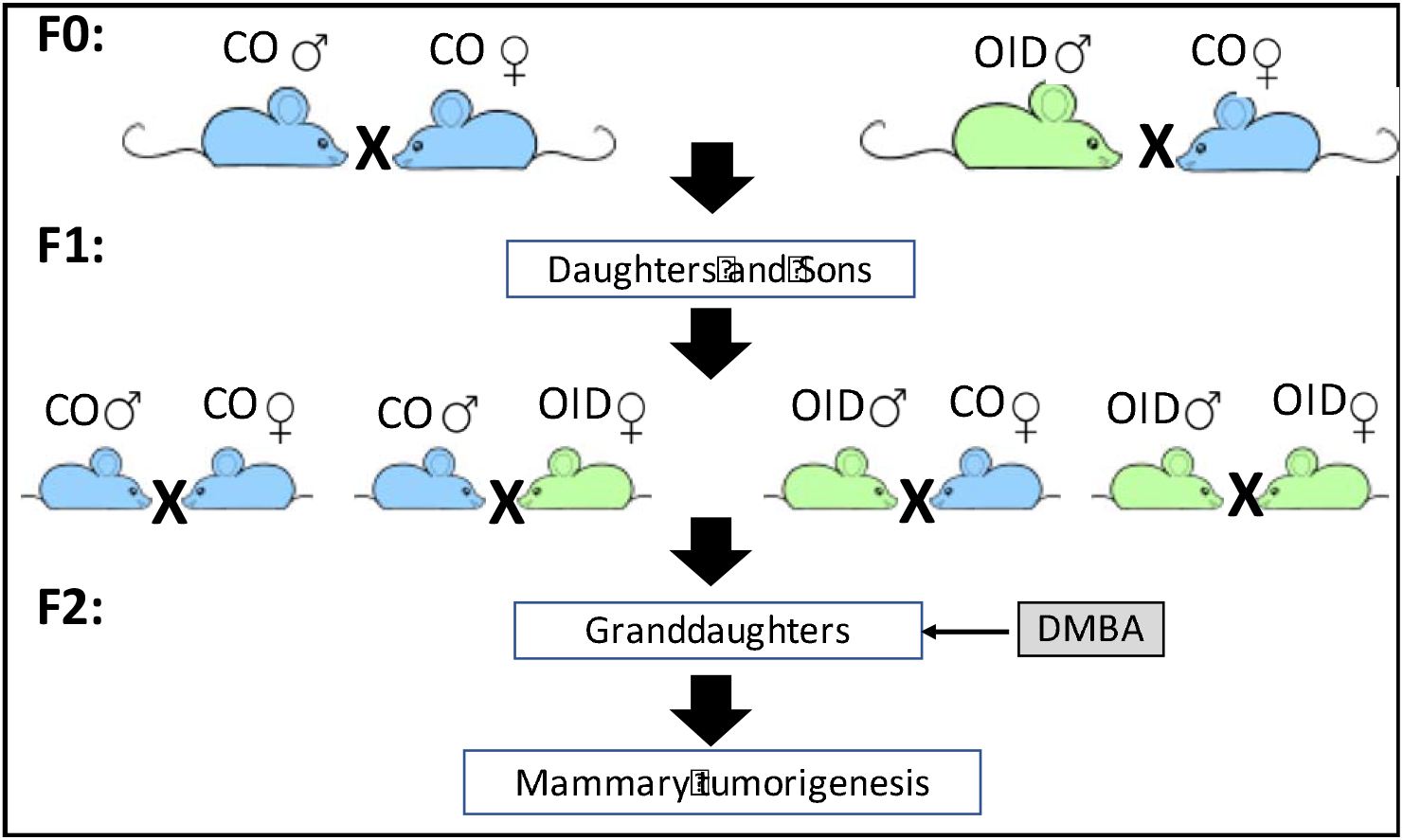
Breeding scheme to produce the F1 and F2 generations: Male mice were fed the experimental diets [control (CO) or obesity-inducing diet (OID)] from 3 to 10 weeks of age. CO diet or an OID diet-fed male mice (**F0**) were mated with female mice that were reared on a CO diet only. The resulting male and female offspring (**F1**) were used to produce the **F2** generation. No sibling mating was carried out.

**Figure S3:**
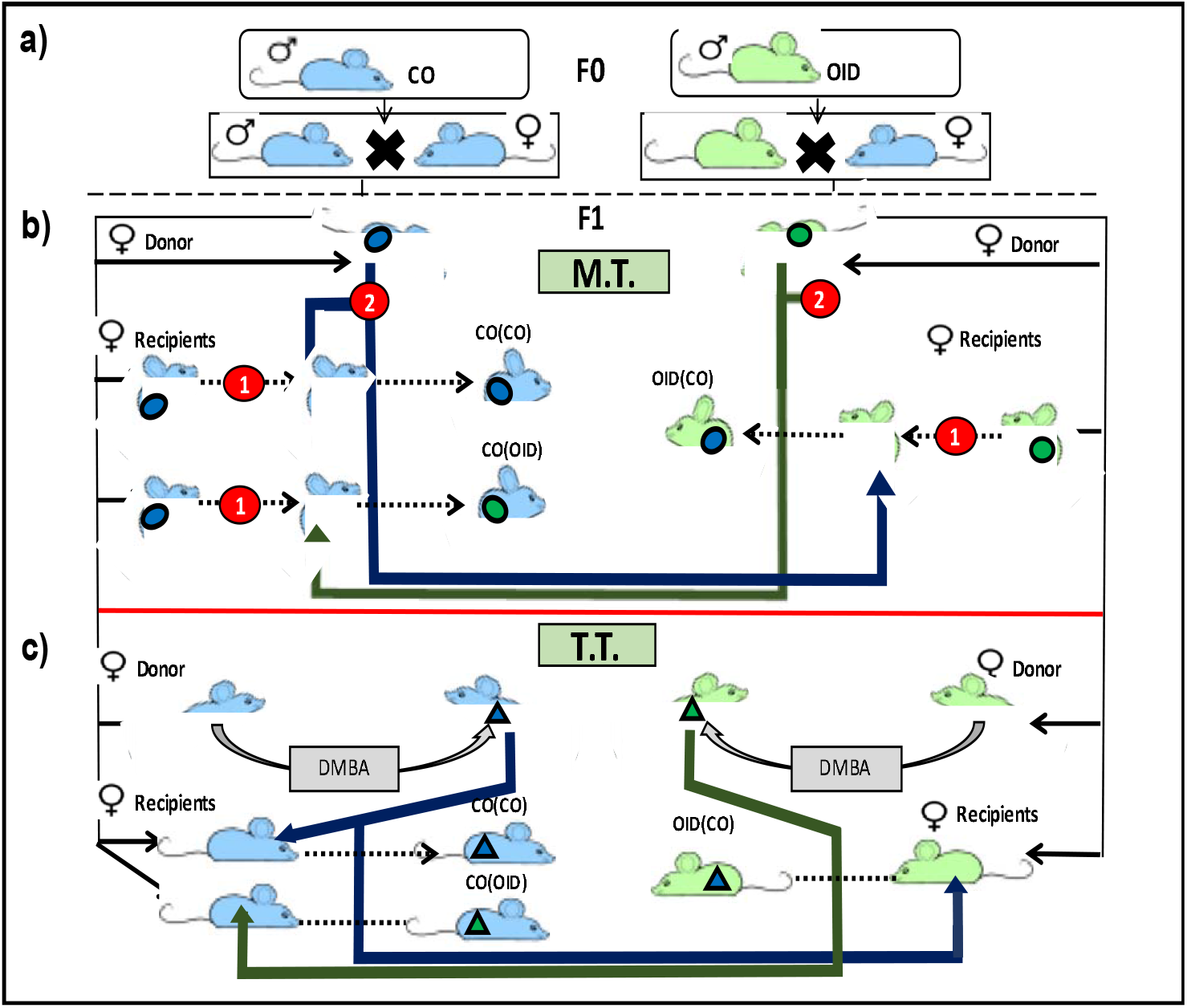
Mammary transplantation study design: **a**) CO diet or an OID diet-fed male mice (F0) were mated with female mice (F0) that were reared on a CO diet only. F1 females, which consumed only CO diet, were submitted to either a **Mammary Transplantation** (M.T.) or to a **Tumor Transplantation** (T.T.). **b**) For the M.T., female recipients (from both CO and OID groups) had their 4th inguinal mammary gland removed (1) and later received a mammary gland transplant (colored circles) (2) from either a donor from the same group or from the opposite group. **c**) For the T.T., female donors received 7,12-dimethylbenz[a]anthracene (DMBA) to induce mammary tumors. Later, female recipients (from both CO and OID groups) received, in their 4th inguinal mammary gland, a tumor transplant (colored triangles) from either a donor from the opposite or from the same group.

**Figure S4:**
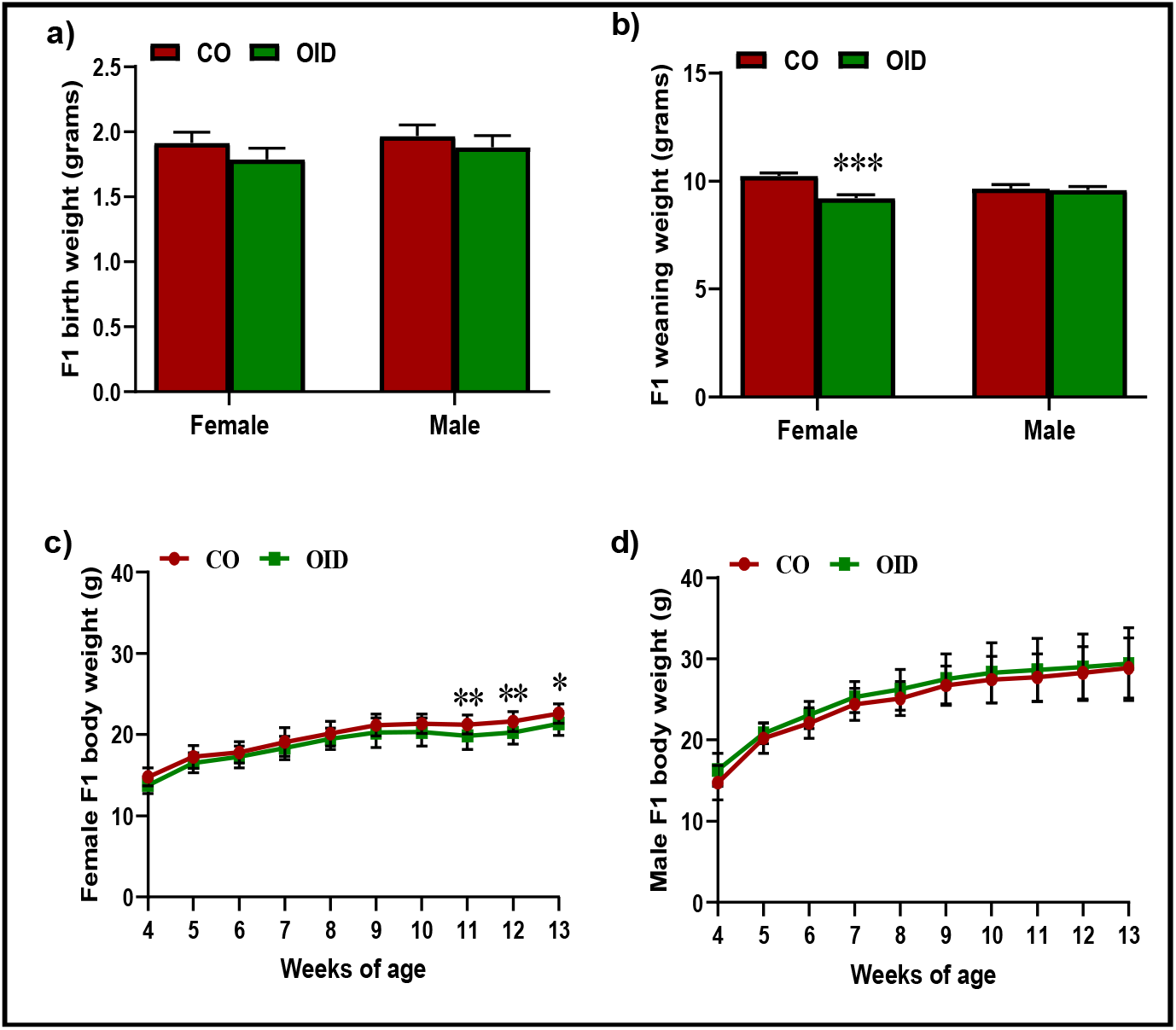
OID and CO F1 generation offspring’s body weight at different stages of life. Birth (**a**), weaning (**b**), and longitudinal body weight in female (n=25/group) (**c**) and male (n=34-43/group) (d) F1 generation offspring from fathers fed with CO and OID diets. The data are expressed as mean ± SEM. Significant differences versus the control group were determined by two-way followed by post-hoc analysis. *P≤0.05; **P≤0.01; ***P≤0.001.

**Figure S5:**
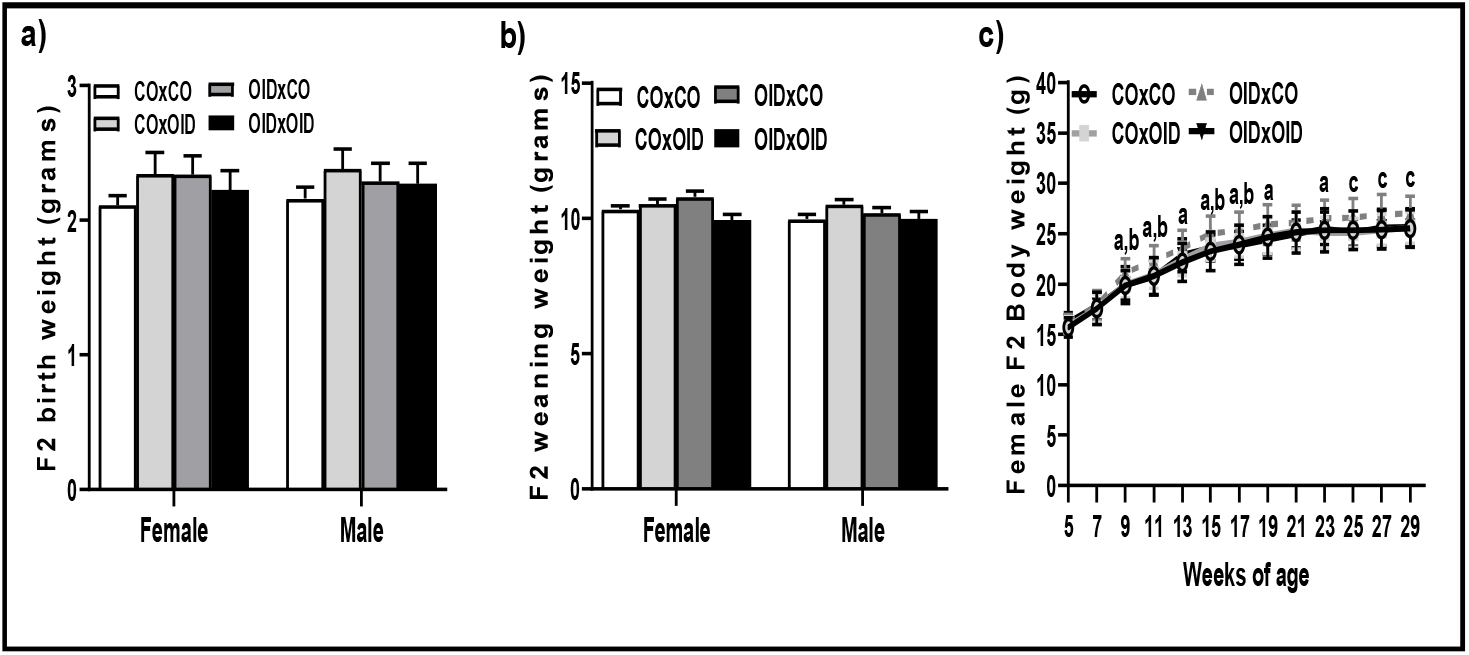
OID and CO F2 generation offspring’s body weight at different stages of life. Birth (**a**)and weaning (**b**) of male and female offspring; Longitudinal body weight (**c**) of female F2 offspring (n=25/group). The data are expressed as mean ± SEM. Significant differences were determined by two-way ANOVA followed by post-hoc analysis. “a” indicates statistically significant difference (P≤0.05) between OIDxCO and COxCO group; “b” indicates statistically significant difference (P≤0.05) between OIDxCO and OIDxOID; “c” indicates statistically significant difference (P≤0.05) between OIDxCO and COxCO, OIDxCO and OIDxOID.

**Table S1:** Composition of the experimental diets

| *Ingredients* | *g/kg* |  |
| --- | --- | --- |
|  | **Control (CO)**<br>**TD.160018** | **Obesity-inducing diet**<br>**(OID)**<br>**TD.160019** |
| Casein | 200.0 | 288.0 |
| L-Cystine | 3.0 | 2.0 |
| Corn Starch | 397.386 | - |
| Maltodextrin | 132.0 | 150.45 |
| Sucrose | 100.0 | 142.0 |
| Corn Oil | 50.0 | 50.0 |
| Lard | 20.0 | 280.0 |
| Cellulose | 50.0 | 20.0 |
| Mineral Mix, AIN-93-MX<br>(94046) | 35.0 | 49.7 |
| Vitamin Mix, AIN-93-VX<br>(94047) | 10.0 | 14.2 |
| Choline Bitartrate | 2.50 | 3.55 |
| TBHQ, antioxidant | 0.014 | - |
| Food Coloring | 0.1 | 0.1 |
| % protein by weight (% **kcal**) | 17.7 (**18.8**) | 25.3 (**19.3**) |
| % carbohydrate by weight (%<br>**kcal**) | 60.1 (**63.9**) | 31.0 (**23.6**) |
| % fat by weight (% **kcal**) | 7.2 (**17.2**) | 33.3 (**57.1**) |
| **Kcal/g** | **3.8** | **5.2** |

**Table S2:** Proportion of female and male offspring in CO and OID litters.

| Generation | Group | Female | Male |
| --- | --- | --- | --- |
| **F1** | **CO** | 0.57±0.2 | 0.43±0.1 |
|  | **OID** | 0.49±0.2 | 0.51±0.2 |
| **F2** | **COxCO** | 0.57±0.5 | 0.43±0.4 |
|  | **COxOID** | 0.45±0.6 | 0.55±0.4 |
|  | **OIDxCO** | 0.52±0.4 | 0.48±0.4 |
|  | **OIDxOID** | 0.56±0.6 | 0.44±0.5 |

All data are mean ± SEM.

**Table S3:** Number of offspring and number of contributing fathers per experiment.

| Samples | Assay | Group | Father | Female |
| --- | --- | --- | --- | --- |
| F1 offspring | Insulin Tolerance Test | CO | 5 | 6 |
|  |  | OID | 4 | 6 |
| Mammary gland transplantation of female F1 offspring | Mammary gland development | CO(CO) | 5 | 5 |
|  |  | CO(OID) | 6 | 9 |
|  |  | OID(CO) | 10 | 12 |
| Mammary tumor transplantation of female F1 offspring | Tumorigenesis | CO(CO) | 7 | 10 |
|  |  | CO(OID) | 7 | 12 |
|  |  | OID(CO) | 6 | 18 |
|  | Ki67 | CO(CO) | 5 | 5 |
|  |  | CO(OID) | 4 | 5 |
|  |  | OID(CO) | 4 | 9 |
|  | Apoptosis | CO(CO) | 4 | 5 |
|  |  | CO(OID) | 3 | 3 |
|  |  | OID(CO) | 6 | 11 |
| F2 offspring | Insulin Tolerance Test | COxCO | 5 | 8 |
|  |  | COxOID | 6 | 8 |
|  |  | OIDxCO | 4 | 8 |
|  |  | OIDxOID | 4 | 8 |
|  | Tumorigenesis | COxCO | 10 | 25 |
|  |  | COxOID | 10 | 25 |
|  |  | OIDxCO | 10 | 25 |
|  |  | OIDxOID | 7 | 25 |

**Table S4:** Mammary gland development in 3-week old female offspring of CO and OID male mice.

| Parameters | CO | OID |
| --- | --- | --- |
| **Epithelial Branching** | 2.0±0.3 | 2.6±0.4 |
| **Number of TEBs** | 5.4±1.0 | 6.0±1.0 |
| **Epithelial Elongation (cm)** | 0.30±0.03 | **0.41±0.03\*** |

All data are mean ± SEM (n=6/group). \*P≤0.05 by t-test.

